## Supplementary figures and images for "A conserved protein tyrosine phosphatase, PTPN-22, functions in diverse developmental processes in *C. elegans*"

### Supplemental Figures 1-6

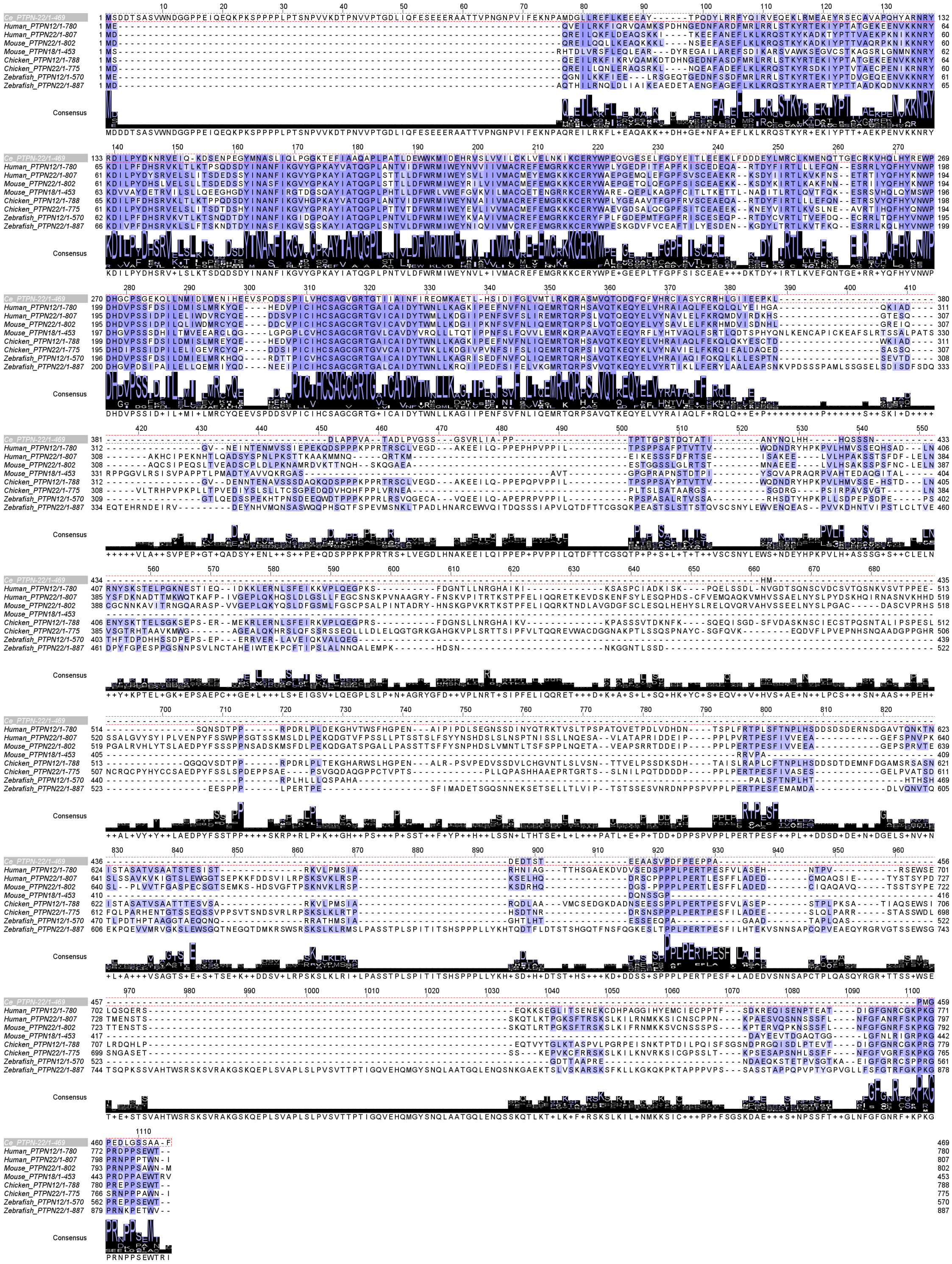

**S2 Fig**

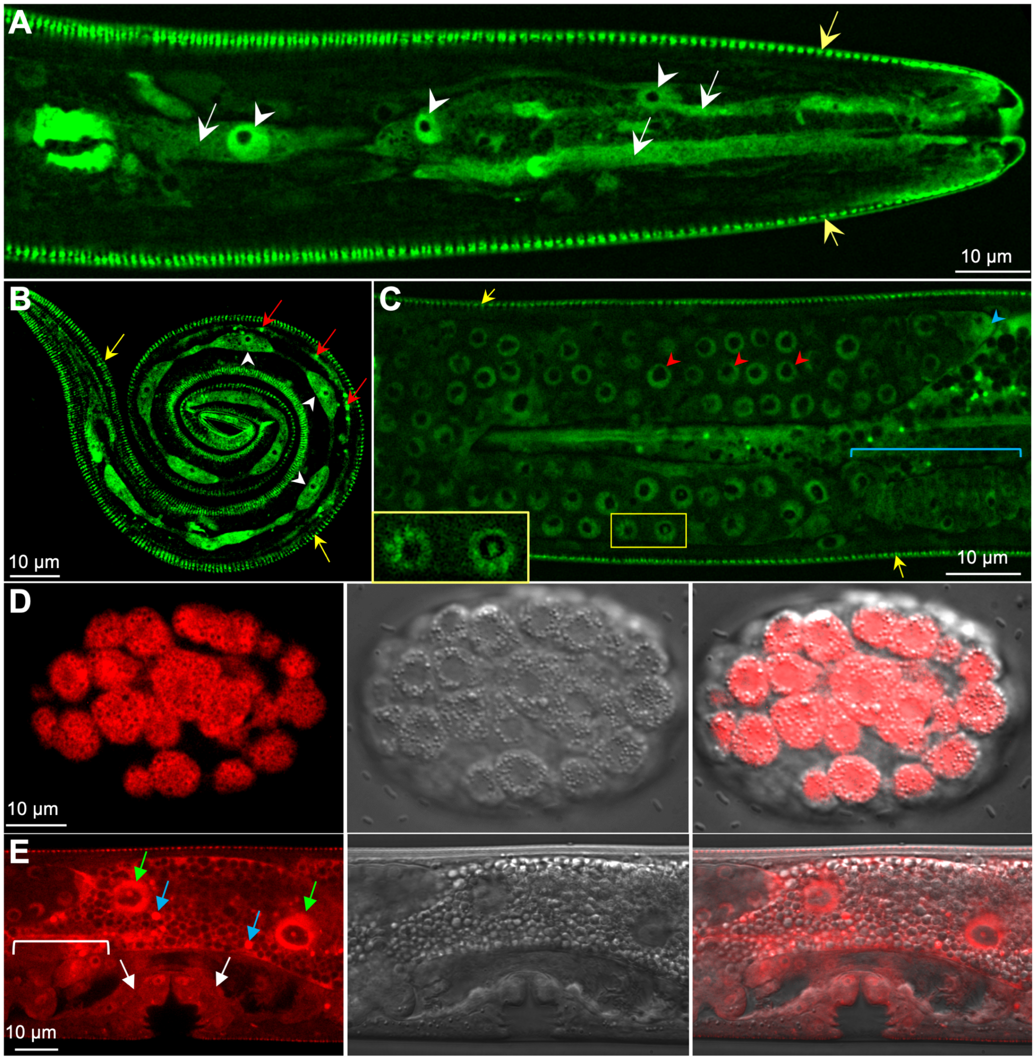

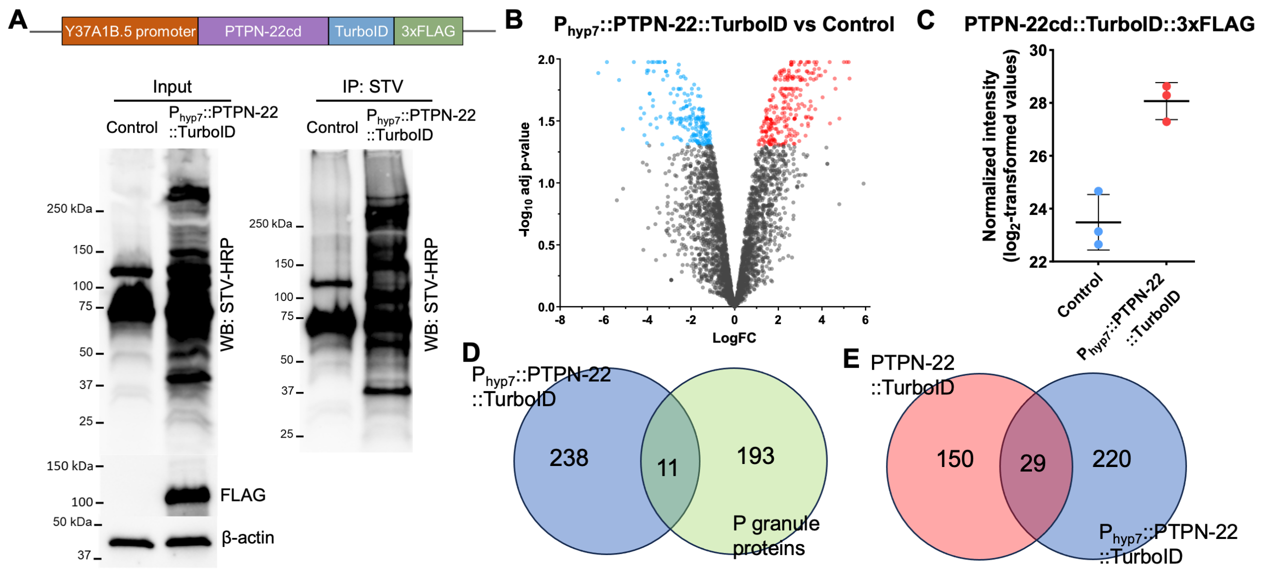

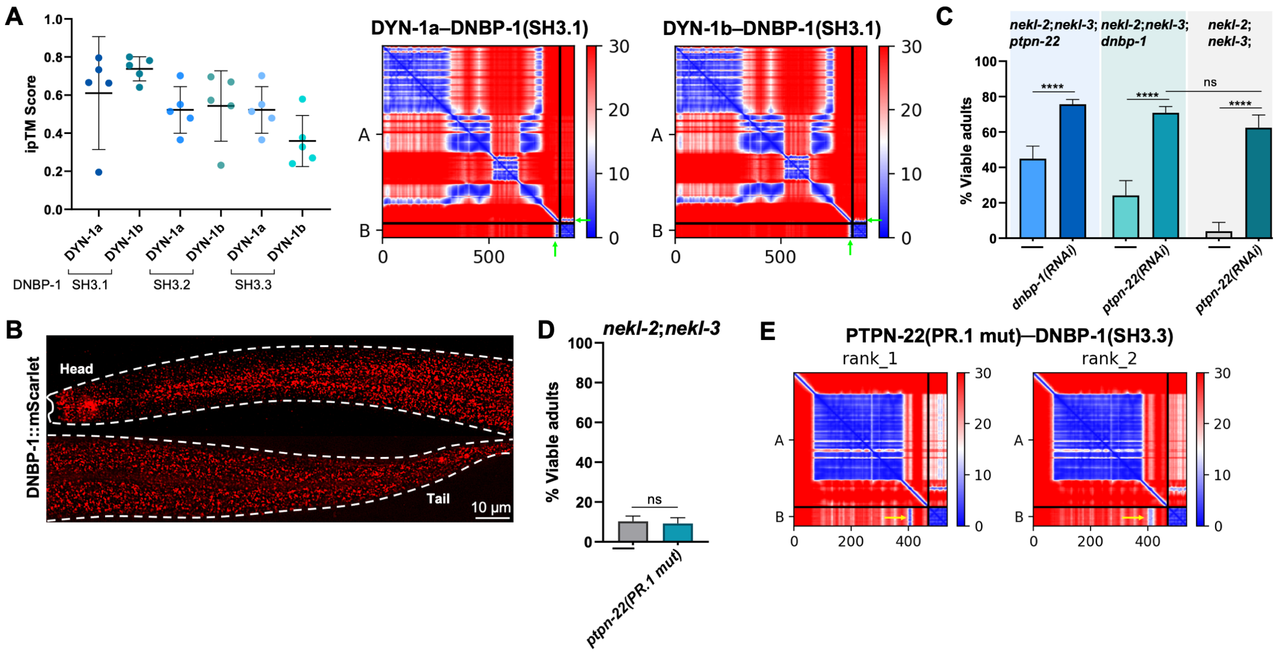

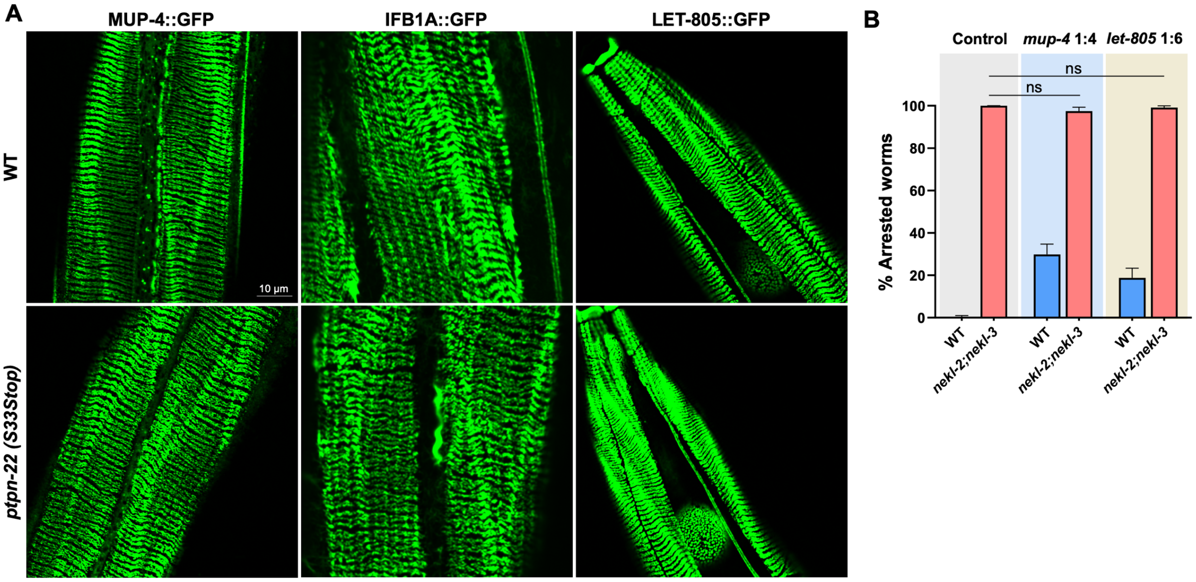

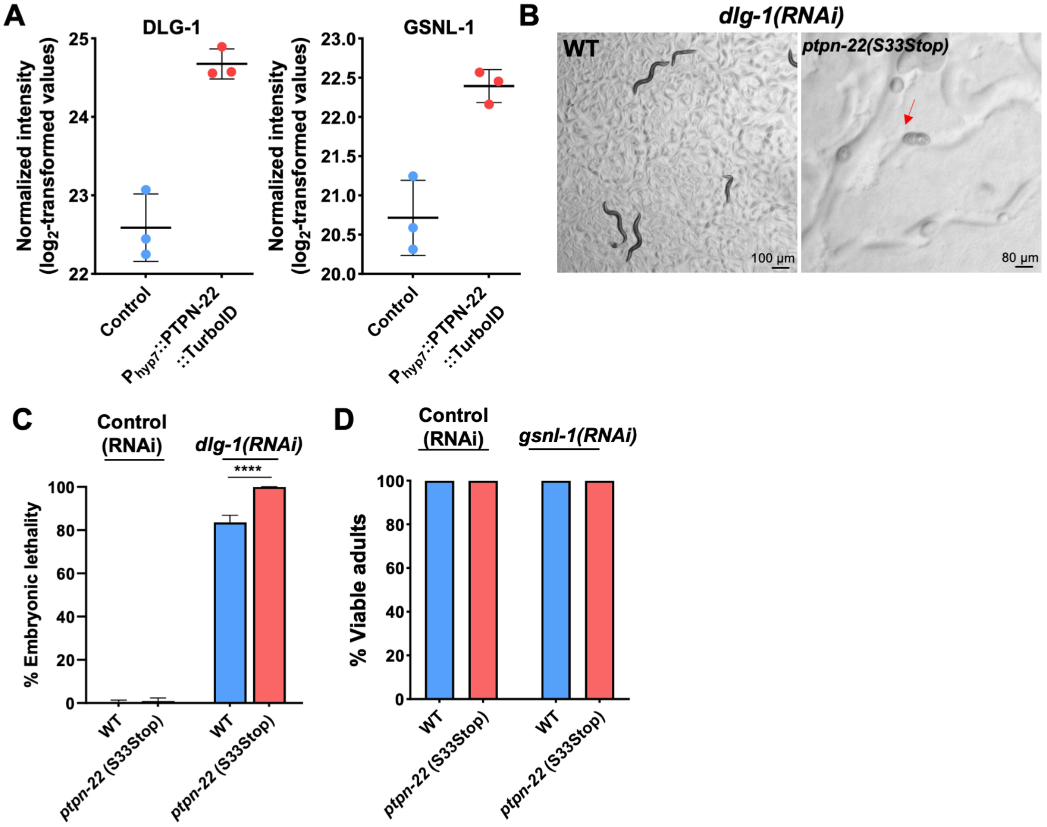
