## Supplementary material for "A conserved protein tyrosine phosphatase, PTPN-22, functions in diverse developmental processes in *C. elegans*": S1 file

**Red = new nucleotide, yellow = mutated nucleotide, * = STOP codon, lowercase letter = intron, capital letter = exon, and green = novel amino acid before the stop codon**

***ptpn-22(fd269)D239fs***

**DNA sequence:**

GGTGATTACGAGATCACTTTAGAAGAGGAGAAACTCTTTGACATGATGTAATATTTAATGAGATGTCTGATGAATATTTAATGA

**Protein sequence:**

673 GGTGATTACGAGATCACTTTAGAAGAGGAGAAACTCTTTGAC 714

225 G D Y E I T L E E E K L F D 238

715 ATGATGTAATATTTAATGAGATGTCTGATGAATATTTAATGA 756

239 M M * Y L M R C L M N I * * 252

***ptpn-22(fd331)E232fs***

**sgRNA:** TTACGAGATCACTTTAGAAG

**Repair template:**
AGAACAAGTCGGTGAAAGTGAACTTTTCGGTGATTACGAGATCACTTTAGACTAGTAGAAACTCTTTGACGATGATGAATATTTAATGAGATGTCTAAAAATGGAGA

**DNA sequence:**

GGTGATTACGAGATCACTTTAAGAAGAGGAGAAACTCTTTGACGATGATGAATATTTAATGAGATGTCTAAAAATGGAGAA

**Protein sequence:**

673 GGTGATTACGAGATCACTTTAAGAAGAGGAGAAACTCTTTGA 714

225 G D Y E I T L R R G E T L * 238

715 CGATGATGAATATTTAATGAGATGTCTAAAAATGGAGAA 753

239 R * * I F N E M S K N G E 251

***dnbp-1(fd385)***

**sgRNA:** ACGTTGCTTCATGTTTTGGA

**Repair template:**

GGTGGACGATGATGATGTTCCAGGTTTTGAGCCTTTTGGAGACACAGAACCATTAACGTTACTA
CATgttttggatttcctggaagaagaacaagcattaagattaataaaaaatattgtttgcaact

**DNA sequence:**
GAAATCCAAGCTTTTGGAGTTGCAAACAATATTTTTATTAATCTTAATGCTTGTTCTTCTTCC
AGGAAATCCAAAACATGTAGTAACGTTAATGGTTCTGTGTCT

**Protein sequence:**

2701 CAGGATCGTATGAAGCCGTTGCAAAAGATTGCGAATAAGGAAATC 2745

901 Q D R M K P L Q K I A N K E I 915

2746 CAAGCTTTTGGAGTTGCAAACAATATTTTTATTAATCTTAATGC 2790

916 Q A F G V A N N I F Y * S * C 930

***dnbp-1(fd386)***

**sgRNA:** ACGTTGCTTCATGTTTTGGA

**Repair template:**

GGTGGACGATGATGATGTTCCAGGTTTTGAGCCTTTTGGAGACACAGAACCATTAACGTTACTA
CATgttttggatttcctggaagaagaacaagcattaagattaataaaaaatattgtttgcaact

**DNA sequence:**
GCCATCATCATCGTCCACCACTGTCACTGAATCACCTTGGGCAACTGCTCC

**Protein sequence:**

2701 CAGGATCGTATGAAGCCGTTGCAAAAGATTGCGAATAAGGCCATC 2745

901 Q D R M K P L Q K I A N K A I 915

2746 ATCATCGTCCACCACTGTCACTGAATCACCTTGGGCAACTGCTCC 2790

916 I I V H H C H * I T L G N C S 930

***ptpn-22(fd388)S33Stop***

**sgRNA:** TTGACTACTGGATTCGATGT

**Repair template:**

tgtttttagcctaaattttcgctattttcgaacCATTTGGAGTATCTTTAGACTACTTAAGTC TATGTATTCAGTGGCGGCGGTGGGCTTTTTGGCTTTTGTTCCTGAATTTCctgaaaaa

**DNA sequence:**

GAAATTCAGGAACAAAAGCCAAAAAGCCCACCGCCGCCACTGAATACATAGACTTAAGTAGTCT AAAGATACTCCAAA

**Protein sequence:**

43 CCTCCAGAAATTCAGGAACAAAAGCCAAAAAGCCCACCGCCG 84

15 P P E I Q E Q K P K S P P P 28

85 CCACTGAATACATAGACTTAAGTAGTCTAAAGATACTCCAAA 126

29 P L N T * T * V V * R Y S K 42

***ptpn-22(fd390)S33Stop***

**sgRNA:** TTGACTACTGGATTCGATGT

**Repair template:**

tgtttttagcctaaattttcgctattttcgaacCATTTGGAGTATCTTTAGACTACTTAAGTC TATGTATTCAGTGGCGGCGGTGGGCTTTTTGGCTTTTGTTCCTGAATTTCctgaaaaa

**DNA sequence:**

GAAATTCAGGAACAAAAGCCAAAAAGCCCACCGCCGCCACTGAATACATAGACTTAAGTAGTCT AAAGATATAATTCG

**Protein sequence:**

43 CCTCCAGAAATTCAGGAACAAAAGCCAAAAAGCCCACCGCCG 84

15 P P E I Q E Q K P K S P P P 28

85 CCACTGAATACATAGACTTAAGTAGTCTAAAGATATAATTCG 126

29 P L N T * T * V V * R Y N S 42

***ptpn-22(fd406)C306S***

**sgRNA:** AATCGGTGAAGAATCTTGGG

**Repair Template:**

GAATAAAATTAATCGCAATAATTGTGCCCGTTCTACCAACACCCGCACTAGAATGTACTAGTAT CGGTGAAGAATCTTGGGGAGACACTTCTTCATGAATATTCTCCATTAAATCAATCATATTCAAT AGTTG

**DNA sequence:**

TGGAGAATATTCATGAAGAAGTGTCTCCCCAAGATTCTTCACCGATACTAGTACATTCTAGTGC GGGTGTTGGTAGAACGGGCACAATTATTGCGATTAATTTTATT

**Protein sequence:**

883 TCTCCCCAAGATTCTTCACCGATACTAGTACATTCTAGTGCG 924

295 S P Q D S S P I L V H S S A 308

925 GGTGTTGGTAGAACGGGCACAATTATTGCGATTAATTTTATT 966

309 G V G R T G T I I A I N F I 322

***ptpn-22(fd444)* Proline-rich region mutated using CRISPR**

**sgRNA:** ATGTAGGCAGTGGCGGCGGT

**Repair template:**

AgcctaaattttcgctattttcgaacCATTTGGAGTATCTTTGACTACTGCATTCGATGTAGCC AGTGCCGCAGCTGCGCTTTTTGGCTTTTGTTCCTGAATTTCctgaaaaaaaccaaattttatg

**DNA sequence:**

GAAATTCAGGAACAAAAGCCAAAAAGCGCAGCTGCGGCACTGGCTACATCGAATGCAGTAGTCA AAGATACTCCAAATGgttcgaaaatagcgaaaattt

**Protein sequence:**

43 CCTCCAGAAATTCAGGAACAAAAGCCAAAAAGCGCAGCTGCG 84

15 P P E I Q E Q K P K S A A A 28

85 GCACTGGCTACATCGAATGCAGTAGTC 111

29 A L A T S N A V V 37

***ptpn-22(syb6505)***

**DNA sequence:**

CCTATGGGGCCGGAAGATCTTGGCAGCAGCGGCATTTGGAGGTGGTGGATCAGGCTCGGGAG
GTCGAGGCTCAGGATCCGGTTCCGGCTCCGGCTCTGGTTCCGGTTCGGGTTCCGGTTCTGGAAA
GGATAACACCGTTCCACTTAAGCTTATCGCCCTTCTTGCCAACGGAGAATTCCACTCTGGAGAG
CAACTTGGAGAGACTCTTGGAATGTCCCGTGCTGCCATCAACAAGCATATCCAAACCCTTCGTG
ATTGGGGAGTTGATGTTTTCACTGTTCCAGGAAAGgtaagtttaaacatatatatactaactaa
ccctgattatttaaattttcagGGATACTCCCTTCCAGAGCCAATCCCACTTCTTAACGCCAAG
CAAATCCTTGGACAACTTGATGGAGGATCCGTCGCTGTCCTTCCAGTTGTTGATTCCACCAACC
AATACCTTCTTGACCGTATCGGAGAGCTTAAGTCTGGAGACGCCTGCATCGCTGAGTACCAACA
AGCTGGACGCGGATCTCGCGGACGCAAGTGGTTCTCCCCATTCGGAGCCAACCTTTACCTTTCT
ATGTTCTGGCGTCTTAAGCGTGGACCAGCTGCTATCGGACTTGGACCAGTTATCGGAATCGTTA
TGGCTGAGGCCCTTCGTAAGCTTGGAGCTGATAAGgtaagtttaaacagttcggtactaactaa
ccatacatatttaaattttcagGTTCGTGTTAAGTGGCCAAACGATCTTTACCTTCAAGACCGT
AAGCTTGCTGGAATCCTTGTCGAGCTTGCTGGAATCACCGGAGACGCCGCTCAAATCGTTATCG
GAGCTGGAATCAACGTTGCCATGCGTCGTGTTGAGGAGTCTGTTGTTAACCAAGGATGGATCAC
TCTTCAAGAGGCTGGAATCAACCTTGATCGTAACACCCTTGCTGCCACCCTTATCCGTGAGCTT
CGTGCTGCCCTTGAGCTTTTCGAGCAAGAGGGACTTGCCCCATACCTTCCACGCTGGGAGAAGC
TTGACAACTTCATCAACCGCCCAGTTAAGCTTATCATCGGAGATAAGGAAATCTTCGGAATCTC
TCGCGGAATCGACAAGCAAGGAGCTCTTCTTCTTGAGCAAGATGGAGTCATTAAGCCATGGATG
GGAGGAGAGATTTCCCTTCGTTCCGCTGAGAAGGCCGGAGGAGATTATAAAGACGATGACGATA
AGCGTGACTACAAGGACGACGACGACAAGCGTGATTACAAGGATGACGATGACAAGTGAatgtg
agaaggcgctttttagcttgaa
